# CCfrag: Scanning folding potential of coiled-coil fragments with AlphaFold

**DOI:** 10.1101/2024.05.24.595610

**Authors:** Mikel Martinez-Goikoetxea

## Abstract

**Motivation:** Coiled coils are a widespread structural motif consisting of multiple α-helices that wind around a central axis to bury their hydrophobic core. Although their backbone can be uniquely described by the Crick parametric equations, these have little practical application in structural prediction, given that most coiled coils in nature feature non-canonical repeats that locally distort their geometry. While AlphaFold has emerged as an effective coiled-coil modeling tool, capable of accurately predicting changes in periodicity and core geometry along coiled-coil stalks, it is not without limitations. These include the generation of spuriously bent models and the inability to effectively model globally non-canonical coiled coils. In an effort to overcome these limitations, we investigated whether dividing full-length sequences into fragments would result in better models.

**Results:** We developed CCfrag to leverage AlphaFold for the piece-wise modeling of coiled coils. The user can create a specification, defined by window size, length of overlap, and oligomerization state, and the program produces the files necessary to run structural predictions with AlphaFold. Then, the structural models and their scores are integrated into a rich per-residue representation defined by sequence-or structure-based features, which can be visualized or employed for further analysis. Our results suggest that removing coiled-coil sequences from their native context can in some case improve the prediction confidence and avoids bent models with spurious contacts. In this paper, we present various use cases of CCfrag, and propose that fragment-based prediction is useful for understanding the properties of long, fibrous coiled coils, by showing local features not seen in full-length models.

**Availability and Implementation:** The program is implemented as a Python module. The code and its documentation are available at https://github.com/Mikel-MG/CCfrag.

## Introduction

Coiled coils consist of multiple α-helices that wind around a central axis to bury their hydrophobic core. They are widespread in proteomes, where they can be found in a variety of forms, depending on the number and orientation of their constituent helices (Lupas and Bassler 2017). This topological diversity is underpinned by the seven-residue heptad repeat, labeled a-g, where the *a* and *d* positions form the core. The geometry of coiled-coil interaction, known as knobs-into-holes, involves the core residue of a helix (knob) packing into a cavity formed by four residues (hole) of an adjacent helix (Crick 1953a). They are the best understood protein fold, as indicated by the number of programs that are able to detect coiled-coil forming propensity from sequence (Lupas, Bassler and Dunin-Horkawicz 2017), and by the existence of the Crick parametric equations that describe their backbone (Crick 1953b). In spite of this, coiled-coil structural prediction has remained a substantial challenge, due to the fact that, although preponderantly repetitive in sequence and structure, coiled-coil domains are rarely without occasional interruptions in the form of non-heptad repeats, which locally alter their packing interactions and geometry.

Even as a general-purpose protein structure model, AlphaFold (Evans *et al*. 2021; Jumper *et al*. 2021) has demonstrated an outstanding ability to accurately model coiled-coil domains, particularly with respect to their supercoiling and core geometry (Madaj *et al*. 2024). Additionally, it has been shown that it can be used to inform of dynamic protein conformations (Wayment-Steele *et al*. 2024; Winski *et al*. 2024). Despite its merits, AlphaFold cannot robustly model some long coiled coils, for which it often outputs oddly bent models that feature spurious contacts or even atomic clashes. We have also observed that it does not generate confident models for non-canonical coiled coils, which are modeled with poor or absent side-chain packing (Martinez-Goikoetxea and Lupas 2023).

Motivated by these observations, we wondered whether computing AlphaFold models of overlapping windows along a sequence would yield better quality models, by simplifying the prediction task. Thus, we developed CCfrag, a pipeline to automate the division of a sequence into fragments and the subsequent integration of the corresponding AlphaFold models into a rich per-residue representation. Our results suggest that not only does this improve the modeling quality of challenging coiled coils, but additionally, it can be used to scan sequences for local structural properties not observed in full-length coiled-coil models. We anticipate that this framework will be of particular interest in the context of understanding long fibrous coiled coils, such as myosins and kinesins.

### Implementation and features

CCfrag is implemented as a Python module that contains two main classes, the divider and the integrator. The former is used to divide a sequence according to a user-defined *specification*, and its output consists of the FASTA files necessary to run the AlphaFold predictions (Fig. 1). After running AlphaFold predictions, the integrator module extracts a number of features from the models, and incorporates them into a rich per-residue representation that can be displayed graphically or used for further analysis. CCfrag also provides limited support for ESMfold (Lin *et al*. 2022), increasing speed at the expense of prediction accuracy.

**Figure 1.**
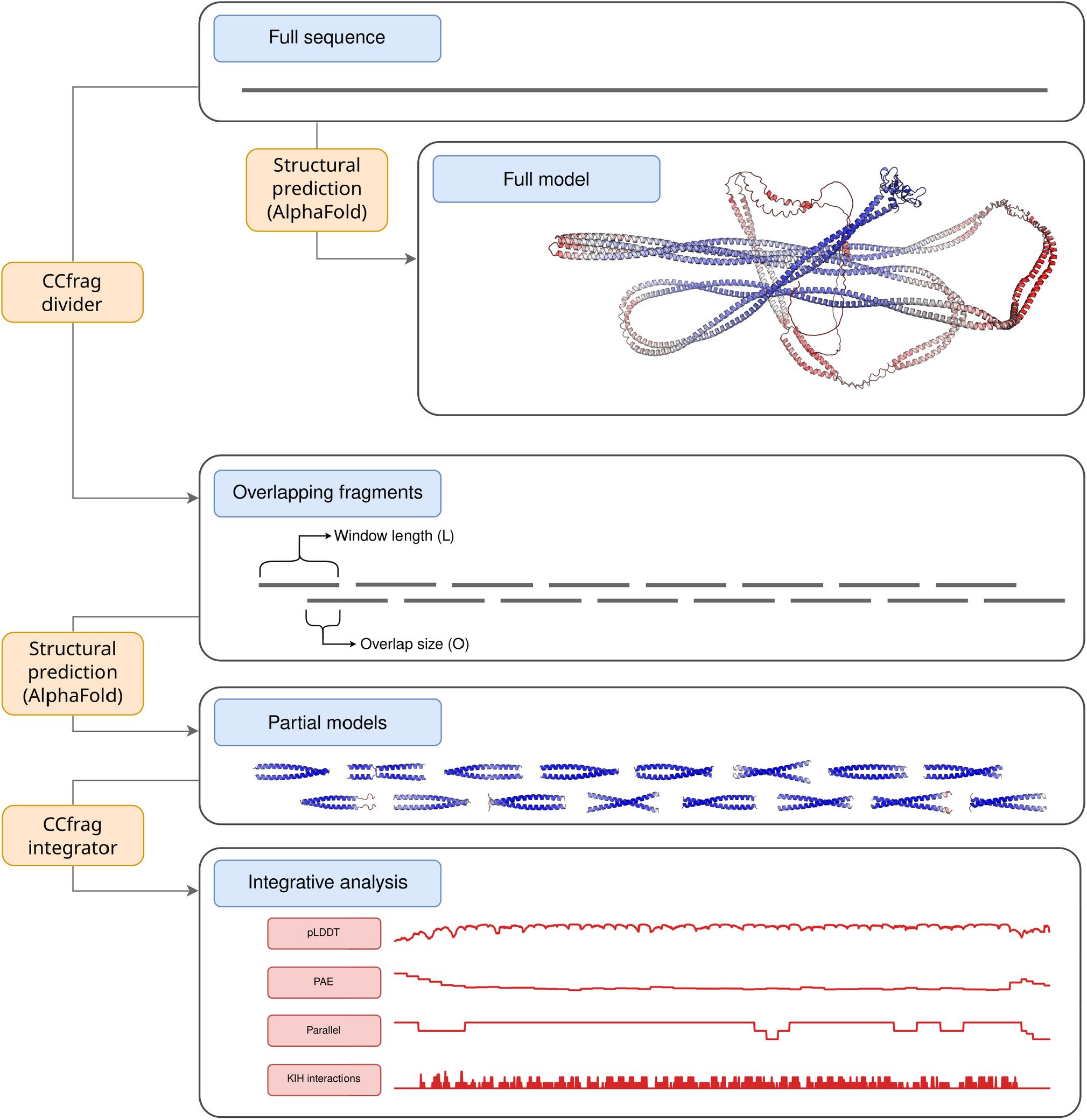
Schematic representation of the CCfrag pipeline. Modeling long coiled-coil domains with AlphaFold generally yields suboptimal models; on top, a full-length model of *H. sapiens* EEA1 is shown, colored by pLDDT (red-worst to blue-best). By dividing the full-length sequence into fragments, the resulting models are predicted with higher confidence, and can be analyzed for local properties not seen in the full-length model (bottom).

The divider module accepts various parameters, the most important of which are the window size (*L*), length of overlap between contiguous fragments (*O*), and oligomeric state (*N*). These define a *specification*, noted in the framework of the program as *N_L*_*O* (for example, 2_30_15 would be 30-residue windows with 15-residue overlap, modeled as a dimer). The minimum overlap size is 0 (no overlap), and the maximum is the length of the sequence minus one. If during the windowing of the input sequence, the C-terminal fragment is not long enough to produce a full window-sized fragment, the window size parameter (*L*) is given priority, and the overlap is increased for that last fragment. An additional feature of CCfrag is the possibility of setting a flanking sequence (*flank*), which will be attached N-and C-terminally to each window for the AlphaFold modeling, but will be removed in the assembly step. The addition of a flanking sequence does not contribute in itself to the scores during assembly (see below), but it can be used to promote the folding of the fragments; this feature is inspired by experimental techniques used to study coiled coils, whereby the addition of GCN4 adaptors is routinely used to promote folding and subsequent crystalization (Hernandez Alvarez *et al*. 2008). The output of the divider module consists of a parameter file (*parameters*.*json*), a table of constructs (*constructs*.*csv*), and a folder (*queries*) which contains the input FASTA files for AlphaFold.

CCfrag does not implement a wrapper to run AlphaFold. This is a necessary limitation given the variety of ways in which users can run AlphaFold (e.g. local machine, high performance cluster), but also a way to decouple the concept of fragment-based modeling from the prediction software itself. Thus, the program can be easily updated to work with newer and potentially better sequence-to-structure prediction programs.

After the AlphaFold models are generated, the integrator module reads most of its required parameters from the configuration file (*parameters*.*json*) and the list of constructs (*constructs*.*cs*v) generated by the divider module. Additionally, the user can specify a list of features that will be extracted or computed from each window. CCfrag includes functions to extract pLDDT and PAE (the average), as well as a function to compute whether the models are parallel or antiparallel, assuming they feature an extended helical conformation. It also includes a wrapper to run SOCKET (Kumar and Woolfson 2021) to detect knobs-into-holes interactions, the hallmark of coiled-coil structures. The addition of arbitrary features (e.g., solvent-accessible surface area) is possible, but requires defining a new function within the source code (an example is provided in the documentation). The output of the integrator module is a table that stores per-residue values for each combination of feature and specification. During the integration process, the features of overlapping windows are flattened via averaging, although this can be customized. This means that, for the case of the parallel/antiparallel feature (numerically encoded as 1 and 0 respectively), some positions of the sequence may have a value of 0.5, meaning that half of the overlapping models showed a parallel arrangement, and the other half an antiparallel one; as illustrated in the next section, this can be interpreted as the lack of topological encoding in the local sequence.

### Case studies

In this section, three examples of the use of CCfrag are presented. The code to generate and visualize these examples is included in the GitHub repository in the form of Jupyter notebooks.

#### Scanning long coiled coils for folding potential

EEA1 (Early Endosome Antigen 1) is a protein that features an N-terminal zinc finger domain, a long parallel dimeric coiled coil, and a C-terminal FYVE domain. It has been extensively studied for its involvement in endosomal trafficking, where its coiled-coil domain is thought to switch between extended and flexible states (Murray *et al*. 2016). Using CCfrag to model EEA1 fragment-wise shows that the pLDDT scores are significantly better than those of the full-length prediction (Fig. 1), with the largest window (L=70) showing the best scores (Fig. S1). On the other hand, it can be observed that shorter windows generally do not encode for a parallel orientation, except for some segments that seem to *pull* from their neighboring residues in larger window sizes; for example, the 400th residue is found in an antiparallel orientation when modeled within a 20-residue context, but the neighboring segments promote the adoption of a parallel arrangement. As expected, there is significant overlap between the SOCKET knobs-into-holes (KIH) detection and the sequence-based coiled-coil predictors DeepCoil2 (Ludwiczak *et al*. 2019) and COILS (Lupas, Van Dyke and Stock 1991). Notably, the absence of KIH interactions also coincides with weak coiled-coil predictions in one or the other program, suggesting that these correspond to flexible regions or segments with ambiguous topological specificity.

#### Scanning for non-canonical coiled coils

In recent work we described a number of protein families that predominantly featured hendecad coiled-coil repeats (Martinez-Goikoetxea and Lupas 2023), and pointed out that these are a particularly difficult targets for AlphaFold. These non-canonical coiled coils are underrepresented in natural proteomes, and possibly as a result, are poorly detected by sequence-based coiled-coil predictors. Using CCfrag to model a member of the MACH family illustrates how piece-wise modeling can be applied to essentially *scan* sequences for potential KIH interactions, thus detecting coiled coils that even coiled-coil specific methods fail to predict (Fig. S2).

#### Multi-state modeling

It has been shown that AlphaFold prediction quality metrics such as pLDDT and PAE can inform of the likely oligomeric states of protein complexes (Madaj *et al*. 2024; Schweke *et al*. 2024). In the context of fragment-based modeling of coiled coils, oligomeric state prediction is particularly challenging due to the fact that coiled-coil sequences can often assemble into various nearly-isoenergetic oligomeric states (Harbury *et al*. 1993). A consequence of this is that when coiled-coil fragments are taken out of their native context, they often crystallize in oligomeric states that do not correspond to the native stoichiometry of the full-length protein. Nevertheless, these fragments inform of the topological specifity that is encoded locally. A great example of this can be found in the spike protein of SARS Coronavirus, whose coiled-coil segments have been crystallized as trimers and tetramers (Deng *et al*. 2006), even though the full-length protein is known to assemble into a trimer. Using CCfrag, we modeled this protein as dimers, trimers, and tetramers, and observed that some of the predicted models matched experimentally-validated coiled-coil domains present in the spike protein (Fig S3). We also observed additional fragments that, even though they do not form coiled coils, were nevertheless predicted as coiled-coil structures. This suggests these fragments feature cryptic coiled-coil folding potentials, and that could fold as such if removed from their native context.

## Conclusions

CCfrag extends the functionality of AlphaFold by facilitating the process of dividing a protein sequence into smaller overlapping fragments, running AlphaFold predictions on these, and integrating the resulting models into a rich representation. We find that this piece-wise modeling can improve the robustness of the predicted models in the case of long coiled coils, even non-canonical ones. Additionally, we propose that this representation can reveal local features not seen in full-sequence models, such as oligomerization preference or folding propensity. CCfrag, together with its documentation and additional examples, is available at https://github.com/Mikel-MG/CCfrag.

## Acknowledgements

We would like to thank Dr Andrei Lupas, Dr Stanislaw Dunin-Horkawicz, Dr Pedro Escudeiro, and Adrian Dobbelstein for useful discussions and comments that contributed greatly to improve this manuscript.

