## Supplemental Figures for "CCfrag: Scanning folding potential of coiled-coil fragments with AlphaFold"

**Supplementary Material**

Mikel Martinez-Goikoetxea

Department of Protein Evolution, Max Planck Institute for Biology, 72076 Tübingen, Germany

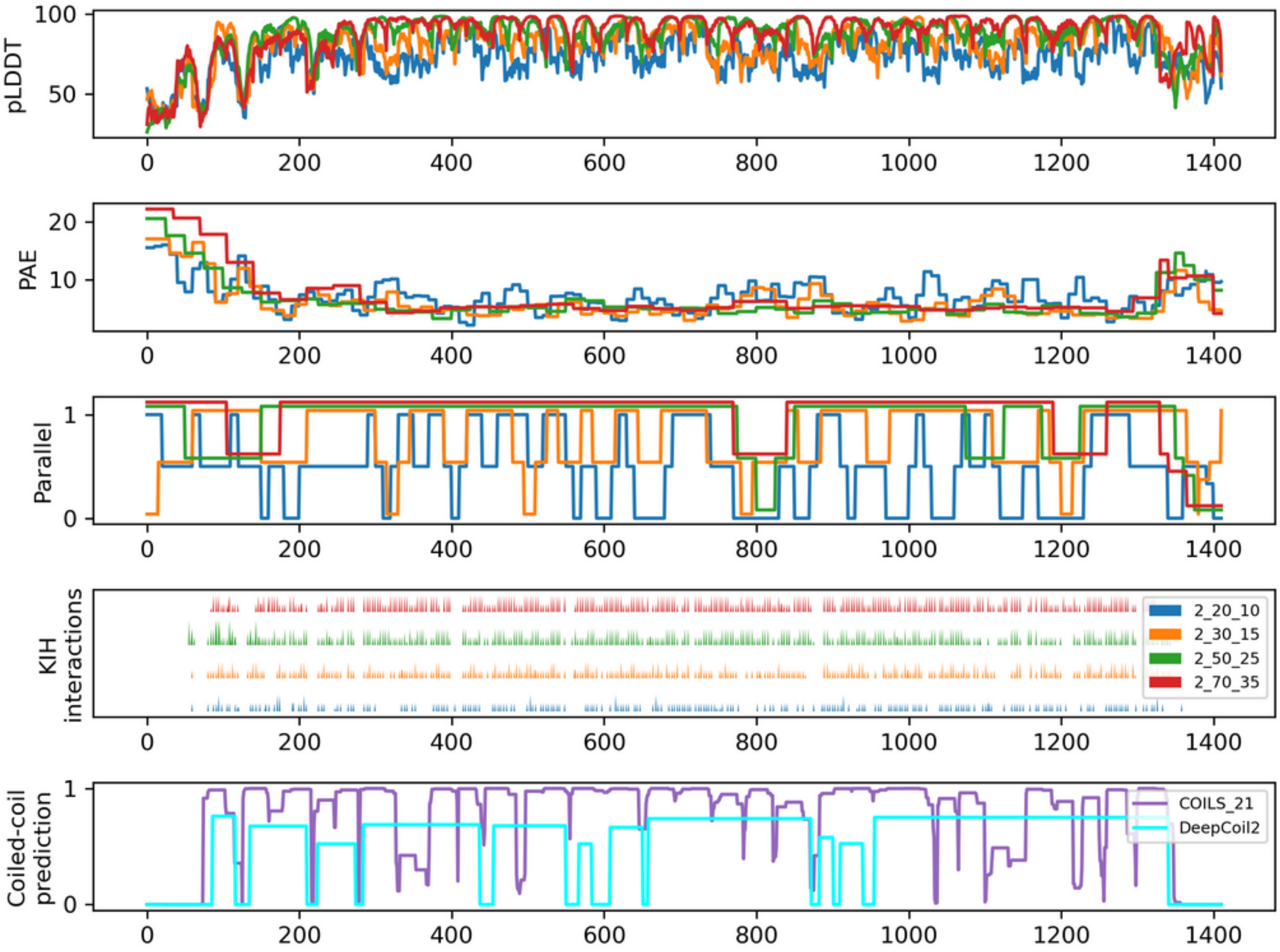


**Figure S1.** Graphical summary of the CCfrag representation of EEA1 of *H. Sapiens*. The protein is modeled as a dimer, in four different specifications with window sizes of 20, 30, 50, and 70 residues and an overlap of half the corresponding window size. In the top 4 panels, plots for various features for each specification (color-coded as in the legend in the right of the fourth panel) are shown (pLDDT, PAE, Parallel, KIH interactions), averaged for each residue (since the overlap is half the window size, each residue is covered by two models, resulting in two values for each feature). In the bottom panel, coiled-coil prediction probabilities for COILS (window size = 21) and DeepCoil 2 are shown.

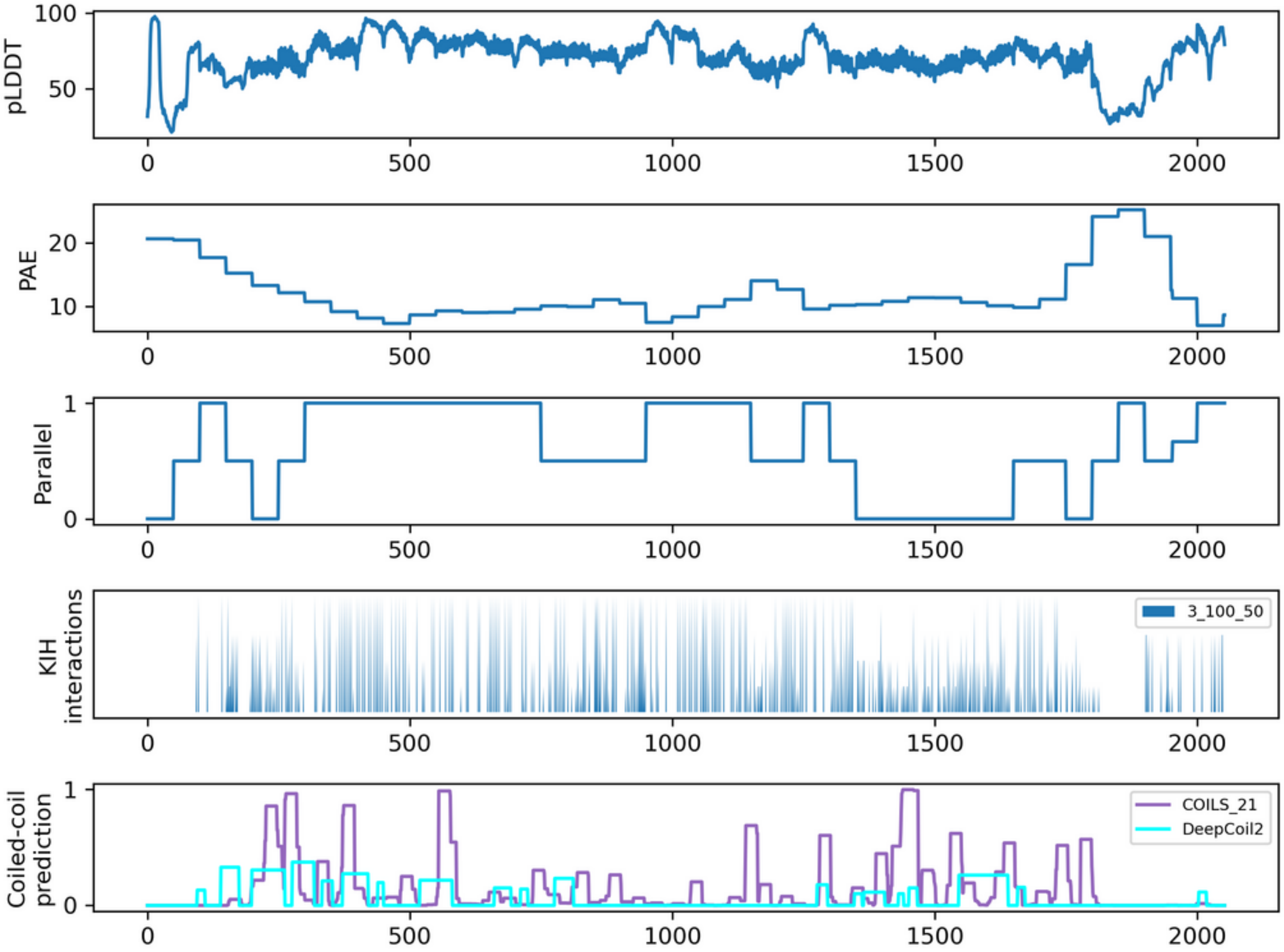


**Figure S2.** Graphical summary of the CCfrag representation of WP_132310275 from *Martelella mediterranea*, a member of the MACH protein family; according to interactive sequence analyses, these proteins feature an extensive hendecad coiled-coil domain. The protein is modeled as a trimer, in a specification of 100-residue windows with 50-residue overlap. Note that even though the sequence-based coiled-coil prediction is very poor, the knobs-into-holes (KIH) interactions can be detected in the structural models along subsequent models, supporting a long coiled-coil stalk.

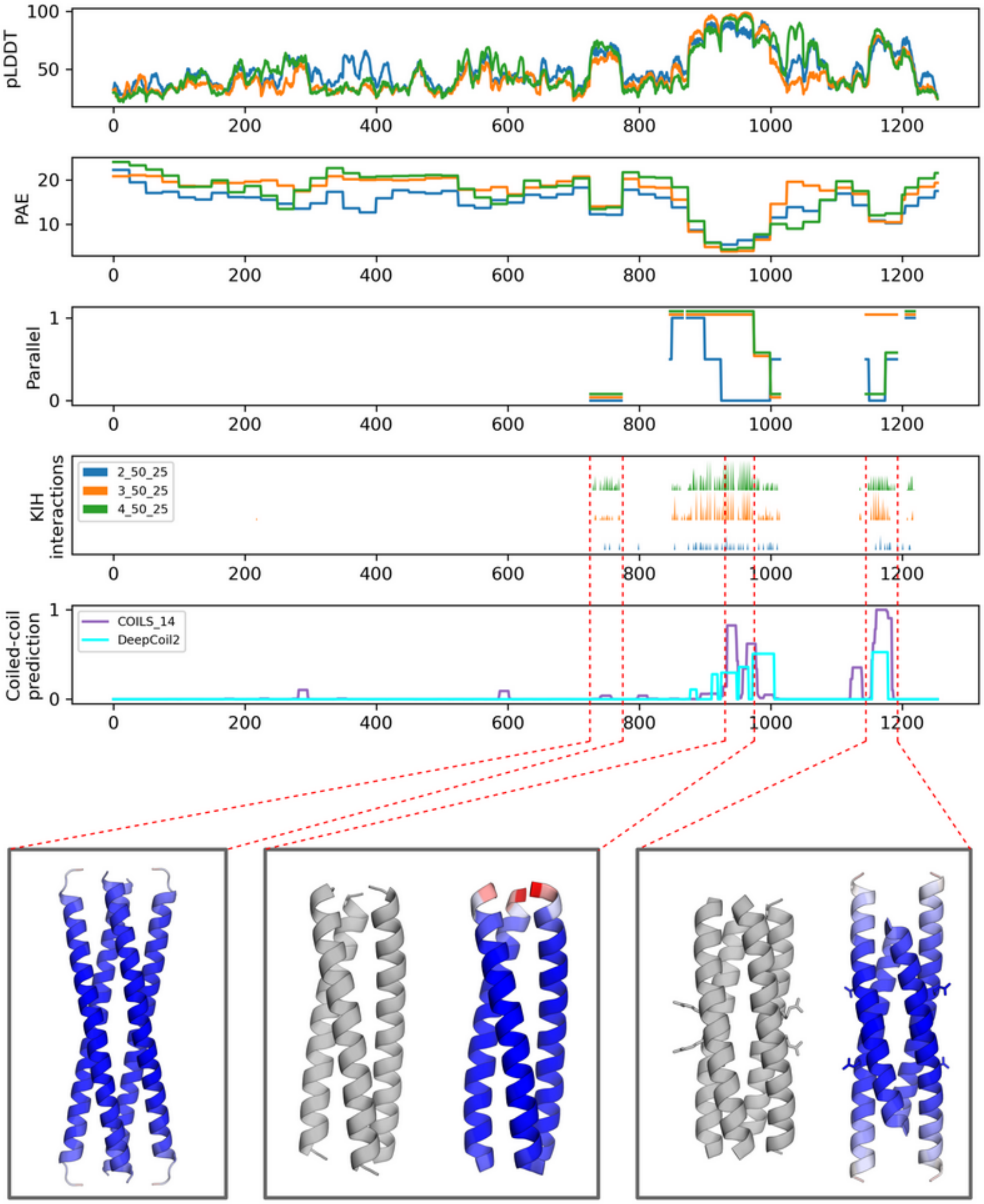


**Figure S3.** Graphical summary of the CCfrag representation of the spike protein of human SARS coronavirus (UniProt P59594). The spike protein is involved in the fusion between the viral and cellular membranes, and although it is not a fibrous protein, it features two functionally relevant coiled-coil domains, known as HR1 and HR2 (for heptad repeat). The protein is modeled as a dimer, a trimer, and a tetramer, each in 50-residue windows with a 25-residue overlap. There are three regions which are more confidently predicted than their surrounding context (high pLDDT, low PAE), two of which correspond to HR1 and HR2. Bottom panel: Left) Fragment 725-775 of the protein is predicted as an interlocking array of coiled coils; Center) Fragment 925-975 is predicted as a parallel trimer, nearly identical to the experimentally solved structure (1ZVB, in grey; RMSD: 0.4 Angstroms). Right) Fragment 1150-1200 is predicted as an antiparallel tetramer, although with a register different of that of the experimentally solved structure (PDB 1ZV7, in grey). The predicted structures are colored by pLDDT (red-worst to blue-best).
